# The interaction between NC(p7)_1-55_ and p6 may regulate interactions with nucleic acids during assembly through modulation of Gag folding

**DOI:** 10.64898/2026.08.28.747767

**Authors:** Valéry Larue, Sylvie Nonin-Lecomte

**Affiliations:** Université Paris Cité, CNRS, CiTCoM, UMR 8038, 4 avenue de l’Observatoire, F-75006 Paris, France

**Author notes:** These two authors contributed equally to the work.

**Keywords:** HIV-1, NC(p7)_1-55_, p6, structure, interaction, assembly, maturation, NMR, Fluorescence, titration, K_d_

## Abstract

We present the solution structures of HIV-1 proteins NC(p7)_1-55_ corresponding to the full-length NC(p7) and mature p6. The studies were carried in water and, to mimic the membrane, in micellar DPC (Dodecylphosphocholine) conditions. Our results unravel for the first time the structure adopted by the N-terminal amino acids of the free NC(p7)_1-55_, with the formation of a small helix spanning residues F6 to R10. Our NMR and Fluorescence Anisotropy data disclose an interaction between NC(p7)_1-55_ and p6 both in water and DPC, with respective K_d_ of 2.5mM and 370 mM at 23°C. The interaction is thus strengthened in lipidic conditions. Protein p6 stabilizes the N-terminus of NC(p7)_1-55_ while increasing at the same time the dynamic of the first zinc finger. Although the entire p6 sequence is involved in the interaction, we show that its C-terminal region is particularly sensitive to the presence of NC(p7)_1-55_, with a propensity of forming a α helix ranging from amino acids S111 to F116. This study brings experimental evidence of a direct protein-protein interaction between p6 and the N-terminal region of NC(p7)_1-55_. We further show that such interaction is readily accommodated within the NC(p15) framework and hypothesize that it may facilitate the selective assembly of assembly of the viral genomic RNA (gRNA) in the cell.

**IMPORTANCE:** Using NMR and Fluorescence Anisotropy spectroscopies, we bring the first experimental demonstration of the formation of α helix in the N-terminus of NC(p7)_1-55_ and of an interaction between the NC(p7)_1-55_ and p6 proteins, both derived from the HIV-1 Gag polyprotein. Such interaction stabilizes the preexisting N-terminal helix while increasing the dynamics of the first zinc finger of NC(p7)_1-55_. The C-terminus of p6 also undergoes conformational changes leading to the formation of a helix. The *in vitro* interaction is strengthened in DPC conditions above the CMC compared to water, suggesting that it could occur *in vivo* in the vicinity of the membrane. The intramolecular interaction between NC(p7) and p6 is readily accommodated in a 3D model of NC(p15). This suggests that p6 could modulate the recognition of the viral genomic RNA (gRNA) by interacting with NC(p7)_1-55_ during assembly and be a key player in this process.

## INTRODUCTION

During the infection by the human immunodeficiency virus (HIV), the Gag protein plays a crucial role in both viral maturation and proliferation stages. Two isoforms, Pr55^Gag^ and Pr160^GagPol^, are expressed at a ratio of approximately 20:1 (1, 2) Pr55^Gag^ is cleaved by the viral protease from its N-terminus to its C-terminus (3) into the Matrice (p17), the Capsid (p24), the SP1 or p2 spacer, the nucleocapsid NC(p7), the SP2 or p1 spacer and the p6 proteins (4)Pr160^GagPol^ cleavage results in additional proteins such as the transframe p6* (TF), the Protease (PR), the Reverse Transcriptase (RT) and the Integrase (IN) (5, 6) Activation of the viral protease and the autocatalytic processing of the precursor polyproteins are essential steps in viral maturation.

Prior to maturation, virion assembly and budding occur in the cytoplasm (7). They are largely driven by interactions between Pr55^Gag^ and the plasma membrane (8), the viral gRNA, as well as with itself (via multimerization) and with other viral proteins. During assembly (8), the MA domain binds to membrane lipids through electrostatic interactions between meristoyl groups at its N-terminus and a neighboring highly basic region (9). MA also interacts with nucleic acids thereby regulating the association between Gag and the cellular membrane. The negatively charged glutamic acid residues in p6 are thought to modulate Gag membrane binding and contribute to the regulation of viral budding (10). The N- and C-terminus domains of CA harbor protein multimerization interfaces that are essential for organizing CA protein-protein interactions into 250 hexameric and 12 pentameric units forming the mature capsid.(11).

The nucleocapsid (NC or NC(p7) _1-55_) and p6 domains are embedded in the C-terminus of Pr55^Gag^ before cleavage (5, 12, 13), separated by the flexible SP2 linker. NC(p7)_1-55_ fulfills multiple critical roles *in vivo* at various stages of the HIV-1 life cycle, in the reverse transcription process (14), the integration of viral DNA (15), the activation of viral RNA dimerization (16), and the facilitation of viral assembly prior to Pr55^Gag^-driven packaging and budding. It exhibits chaperone activity in association with nucleic acids (14, 17). There is general consensus that the nucleocapsid, in its various forms and cellular functions, can either stabilize or destabilize nucleic acid interactions (17), making it a promising target for antiviral strategies (18).

During the assembly process, Pr55^Gag^ was described to interact with the viral genomic RNA (gRNA) and regulate the binding (19). Two copies of unspliced gRNA dimerize in the cytoplasm through their DIS signals (20). The NC(p7)_1-55_ domain contributes to the cytoplasmic clustering of the two strands (21) by interacting with the packaging signals (Psi) located within the gRNA 5ʹ-untranslated region (UTR) and upstream the *gag* gene. In contrast to tRNA^Lys3^, which promotes the formation of a RNA kissing complex during the initiation of the reverse transcription, the nucleocapsid was showed to favor the association of two extended gRNA strands, both in the presence and in the absence of tRNA^Lys3^ (22). Dubois *et al*. (23) have demonstrated that the p6 domain is essential for the specific binding of Pr55^Gag^ to the viral gRNA and that its deletion increases the affinity of Pr55^Gag^ for both cellular RNAs and mutant viral gRNAs. A recent study suggests that, within the context of NC(p15) and NC(p9), p6 domain folds back on NC(p7) thus reducing its NA aggregation capabilities (24, 25).

During the budding phase, the p6 domain interacts with multiple host factors such as the ESCRT associated proteins TSG101 and AIP1/ALIX (26) and with the viral accessory protein Vpr (27) (Fig. 1). TSG101 is a component of the cellular ESCRT-I complex and ALIX of the ESCRT-III complex. Both are key-players in the HIV-1 budding process. In addition to their interaction with p6, the recruitment of TSG101 and ALIX also involves binding to the NC(p7) _1-55_ domain, and a functional interplay between all these components during viral particle budding and release has been proposed (28, 29). Vpr indirectly enhances the efficiency of the viral replication by inducing a G2-arrest of the cellular cycle (30) and promoting the viral pre-integration complex to enter the nucleus of the non-dividing cells (31). It also enhances the viral cytopathogenicity (32, 33). The incorporation of Vpr into the virion requires a direct interaction with the p6 domain of Pr55^Gag^(34).

**Figure 1.**
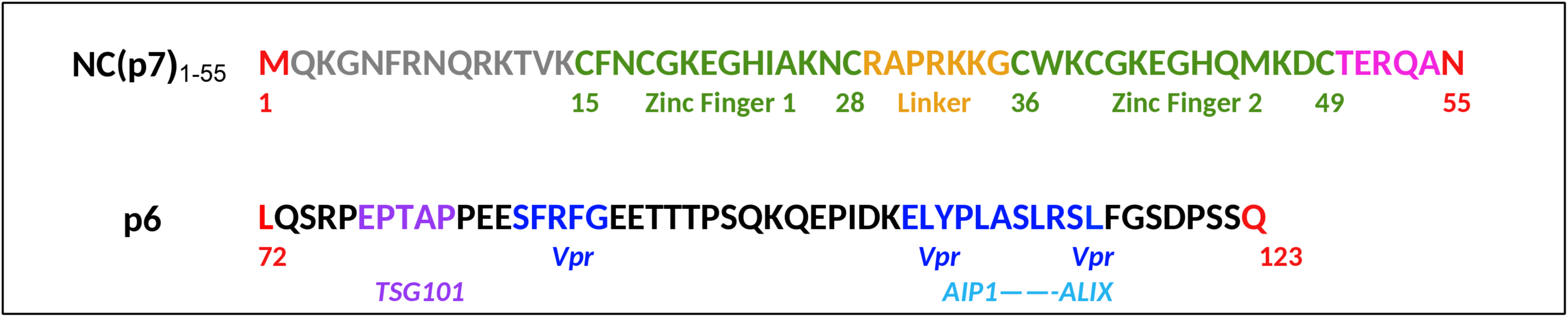
Primary sequences: (**Top)** of NC(p7)_1-55_: N-and C-termini of the protein are in gray and magenta, respectively. The two zinc fingers ZF1 and ZF2 are in green and the basic linker in between in orange. (**Bottom)** p6 protein: the interactions motifs with others VIH partners in infected cell appear in purple and blue.

Because literature suggests the possibility of an interaction between NC(p7)_1-55_ and p6, during the assembling and budding, we have decided to study the structure, the dynamics and the interaction of the two domains, both in aqueous and micellar conditions to mimic the proximity of the membrane. The nucleocapsid structure has been extensively studied. The multiplicity of its functions in the viral cycle justifies the use of diverse environmental conditions observed in the literature for structural studies (*e.g.* salt concentration (35), pH, and solvent (36)). NC(p7)_1-55_ is a metalloprotein which harbors two Zinc Finger motifs (ZF) of the Cys-X_2_-Cys-X_4_-His-X_4_-Cys type, each coordinating one Zn ion with a binding constant of 10^13^-10^14^ M^-1^ (37). These motifs are separated by a linker of 7 amino acids. The ZF formation induces conformational rearrangements that bring functionally distinct aromatic amino acids into proximity (38, 39). The ZF regions are essential for viral infectivity (40). Three-dimensional solution structures of NC(p7) have been solved by nuclear magnetic resonance (NMR). The isolate full-length NC(p7)_1–55_ was described as flexible conformations with no particular structure at the extremities (41, 42). The low structural organization of the whole protein, with large distances spanning the two zinc fingers, was confirmed by RDC NMR and SAXS (43). Residues K3 to K11 form a 3_10_ helix upon interaction with the viral RNA SL2 and SL3 regions (44, 45). Conversely, NC(p7)_13–55_, the truncated version at the N-terminus of NC(p7)_1-55_, exhibits a folded and compact conformation (39, 46), with the aromatic residues of the two ZF spatially close to each other, and with F16 and W37 forming a stacking platform that facilitates interaction with the RNA guanine bases (47). Dynamic analyses based on NMR relaxation data have also been conducted (48). Altogether, these findings demonstrate that the nucleocapsid, free or as part of Pr55^Gag^, adopts a dynamic, sequence- and partner-dependent structure. In solvents containing TFE, p6 adopts a two helical shape conformation (27, 49, 50) whereas in aqueous solution or when embedded within the larger Pr55^Gag^ protein (p6^Gag^), it appears less structured (25). The two helical motifs called late motif I (P-T-A-P) (51) and late motif II (L-Y-P-X) (52) are involved in recruiting host proteins TSG101 (53) and ALIX/AIP1 (52, 54). Additionally, two other helices, helix I (F-X-F-G) (49) and helix II (L-Y-P-X-(n)-L-X-X-L-F-G) (55), have been shown to interact with Vpr (34). Recently, Meshri and collaborators provided evidence that the two zinc fingers of NC(p7) act in concert with p6 to recruit TSG101 in the cytoplasm (53), raising the possibility of a direct interaction between the NC and p6 domains.

In this study, we first present the solution structures of free HIV-1 NC(p7)_1-55_ and p6. We have used NMR spectroscopy in H_2_O and in DPC buffers at 10°C and 30°C and molecular modeling under NMR constraints (angles and distances). We bring experimental evidence that the NC(p7)_1-55_ N-terminus is partially structured and of a direct interaction between regions of NC(p7)_1-55_ and p6, using NMR and Fluorescence Anisotropy spectroscopies. We show that the interaction between NC(p7)_1-55_ and p6 is medium but enhanced in micellar conditions. Our NMR dynamics investigations show that p6 exerts both stabilizing and destabilizing effects on NC(p7). We inject our experimental data to a 3D model of structure of NC(p15) generated by AlphaFold3, and show that the intramolecular interaction between the two domains of Gag is readily accommodated, supporting the idea that p6 may act as one key player for the folding of the NC domain and the regulation of the interaction with the RNA during virion assembly. To assess the potential competition between the viral RNA and p6 for binding to NC(p7)_1-55_, we finally compared the structures of NC(p7) in complex with SL2 (44) and SL3 (45) RNA hairpins to our three-dimensional models of the intramolecular interaction in the context of NC(p15).

## RESULTS

### Structure and dynamic of protein NC(p7)_1-55_ in water and in DPC*-d_38_*

The first additional information brought by our work compared to previous NMR studies, is structural results about the N-terminus of NC(p7)_1-55_ which encompasses 13 residues (Fig. 1). Except for the terminal M1 and Q2 amino acids, the spectral resolution at 600 MHz allowed complete assignment of the 3D [^1^H–^15^N] NOESY-HSQC and 3D [^1^H–^15^N] TOCSY-HSQC spectra of the ^15^N-labelled NC(p7)_1-55_ (hereafter named as ^15^N-NC(p7)_1-55_) in water and at 10°C (Fig. 2A). To assess the impact of the temperature, we performed [^1^H–^15^N] TROSY experiments at 10 and 30°C (Fig. S1A). We observe that, for most of the terminal amino acids prior to ZF1, as well as for R29 and R32 of the linker and G43 and Q53 at the C-terminus, the correlation cross-peaks disappear at 30°C, consistently with either an intermediate exchange regime on the NMR time scale or the destruction of any structural pattern. To explore the impact of a micellar solvent on the dynamics and the structure of the free protein, we also studied ^15^N-NC(p7)_1-55_ in DPC*-d_38_* solvent (450 molar equivalents added), *ie* well above the Critical Micellar Concentration (CMC = 1-2 mM). Except for the N-ter residues, the cross-peak pattern of the NOESY spectrum is very similar in water and in DPC*-d_38_* (not shown). Nevertheless, like observed in water at 30°C, the correlation peaks of amino acids K3 to R10 (N-terminal region), of R29 (linker between the two zinc fingers), and of G43 and Q53 in the C-terminal region, totally disappear from the [^1^H–^15^N] SOFAST-HMQC spectra recorded at 10°C in DPC*-d_38_* (Fig. 2A). They are still broadened or missing at 30°C (Fig. S1B).

**Figure 2.**
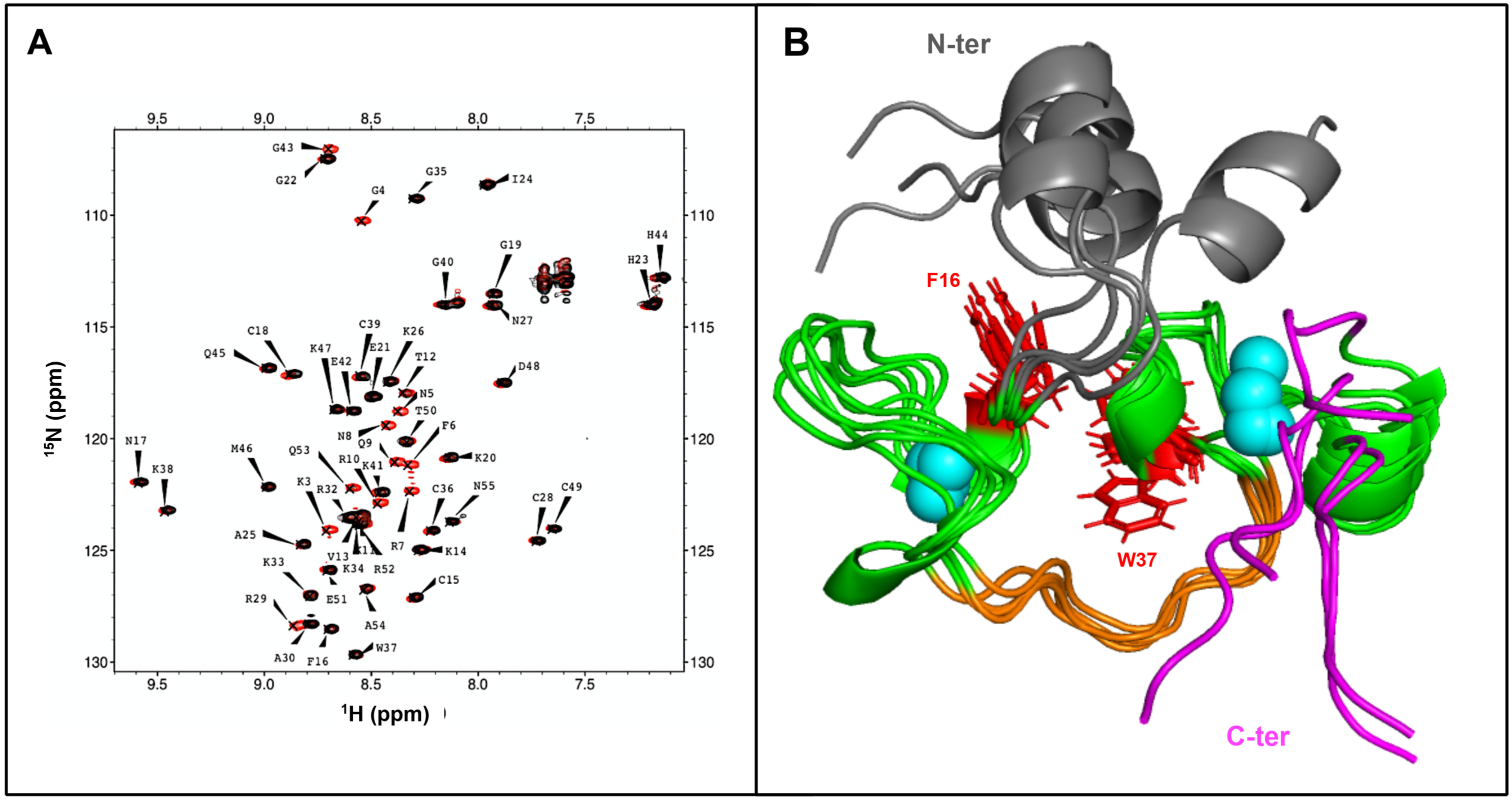
**(A)** Superposition of [^1^H-^15^N] SOFAST-HMQC spectra at 10°C of ^15^N-NC(p7)_1-55_ recorded in H_2_O (red) and with 450 eq. DPC-*d_38_* (black). The peaks of most N-terminal amino acids disappear upon a temperature increase of 20°C. **(B)** Superpositions of the 5 best structural models of free NC(p7)_1-55_ obtained with Xplor-NIH (EEFx force field) softwares with restraints measured at 10°C in water. The N-ter part is gray, the two zinc fingers green, the linker orange and the C-ter part magenta. Aromatic residues F16 and W37 are colored in red. The link between ZF2 and the N-terminal helix (also in gray) is flexible. The N-ter helix adopts different orientations all consistent with the NMR data.

As we wanted to get structural information on the N-ter part of NC(p7)_1-55_, we chose to solve the structure in H_2_O at 10°C and pH 6.5. Interestingly, all our experimental data point to the formation of a helical structure in the N-terminal region. The NMR ^3^J_HN-Hα_ coupling constants obtained from the 3D [^1^H–^15^N] HNHA experiment range from 5 to 7.5 Hz for amino acids N5 to R10, indicating φ angles between -65° and -80° consistent with helical structure (of the α or 3_10_ type). The corresponding ^3^J_N-Hβ_ coupling constants, derived from the 3D [^1^H–^15^N] HNHB experiment, are between 3 and 4.2 Hz corresponding to χ1 angles of -100° to -90°. Both Dangle (Fig. S2) and Talos (not shown) softwares, which use variations in chemical shifts relative to the corresponding random coil values as well as CSI values, support α helical conformation. However, distances restraints derived for NOEs are sparce, like observed previously at 10°C for the N-ter region of NC(p15), probably due to multiple orientations in equilibrium in the intermediate exchange regime. The NMR-constrained molecular models computed with EEFx force field (Fig. 2B) are consistent with all our experimental data. The overall solution structure is organized around two zinc fingers like for the shorter structure previously published (56). It discloses the formation of α helix within the N-ter flexible segment, ranging from N5 to R10, which exhibits two predominant orientations with respect to the first zinc finger depending on 3D models. The atomic coordinates have been deposited in the Protein Data Bank (PDB) under accession code **30ZZ.**

^15^N-H relaxation time measurements were conducted in the presence and absence of DPC-*d_38_* at 10°C (Fig. S3). For residues K11 to N55, the overall aspects of the T1, T2 and ^1^H–^15^N heteronuclear NOEs variations along the sequence are very similar whether in the presence or absence of DPC. T1 values are about 600 ms in water. Only a slight decrease by about 10% is observed in the presence of DPC. T2 relaxation times are nearly identical and around 120 ms in H_2_O and in DPC*-d_38_*. In water, ^15^N-H heteonuclear NOEs undergo a drastic fall starting from V13 and E51 as we move towards the terminal residues Q2 and N55 respectively, disclosing the dynamics of these regions. This also shows that the N-ter helix is embedded in a dynamic region. In DPC*-d_38_*, we could not get access to the relaxation values of the cross-peaks which were broadened to baseline. Fortunately, we could monitor the T1 and T2 variations of R10, which is located at the end of the N-terminal helix according to the 3D models derived from data acquired in H_2_O without any added DPC. R10 T1 and T2 relaxation times are decreased by the presence of DPC*-d_38_*to reach a value comparable to the average of the values observed for the residues embedded in a structure, suggesting a strengthening of the helix. These results collectively evidence that the presence of DPC micelles influences the dynamics of the N-terminal region of NC(p7)_1-55_. We hypothesize that interactions with DPC micelles influences, and most probably slows down, the dynamics of both the N-terminus and the helix, switching from a fast to an intermediate conformational exchange regime that may account for the disappearance of the corresponding NMR signals.

### Structure and dynamic of p6 in water and in DPC*-d_38_*

The 2D ^1^H-^1^H NOESY spectra were recorded with unlabeled p6 in water in the absence or in the presence of added DPC-*d_38_*to study the behavior that p6 might have close to the cell membrane (Fig. 3). Full NH and Hα assignment and almost complete assignment of the lateral chains could be achieved both at 10°C in water and 30°C in the presence of 450 molar equivalents of DPC-*d_38_*. On the 2D NOESY spectra, the low dispersion of the cross-peaks in the NH-NH and NH-Hα regions observed in H_2_O reveals a poor structuring, as already observed in a previous study where p6 is a part of NC(p15) protein (25). Upon addition of DPC-*d_38_* and at 30°C, the cross-peak pattern is different (Fig. 3B). The dispersion is still poor, albeit a little bit better than in water, and the NH-NH cross peaks are more numerous and stronger, supporting a slightly better structuration in the presence of DPC micelles. Chemical shift perturbations are observed upon addition of 450 eq. DPC-*d_38_*. They are mainly clustered in the C-ter part of p6 (Fig S4C). Based on our protons and nitrogens chemical shifts, TALOS predicts more helices in DPC than in water (Fig. S4 A and B), with respectively four (E83 to R87, E91 to T92, I102 to L106, L109 to R113) *vs.* six small helices (E83 to F88, E90 to T92, S96 to K98, K104 to L106, A110 to L115, S121-S122). 3D models of the solution structures of p6 using NMR angle and distance restraints were obtained (Fig. 4). The structure calculations were performed using a set of 155 distance and 80 angle restraints measured in water (Fig. 4A) and a set of 307 distance and 98 angle restraints in DPC-*d_38_* (Fig. 4B). In all for cases, the helical regions are short and discontinuous, in contrast to previously published models of p6 in DPC, which exhibit two major α-helices (27, 50). The DPC models reveal more helical elements than those observed in water, and the helices number and location are close to the helices predicted by TALOS from the chemical shifts (Fig. S4 A and B). Comparison of the structures obtained in water and in DPC reveals increased structuring of the central and C-terminal regions in the presence of DPC (Fig. S4D), which correspond to the regions where NH peaks in the HSQC spectra exhibit the largest CSPs between the two solvents (Fig. S4C). The atomic coordinates have been deposited in the Protein Data Bank (PDB) under accession codes 31AD and 31AA respectively in water and in DPC-d38.

**Figure 3.**
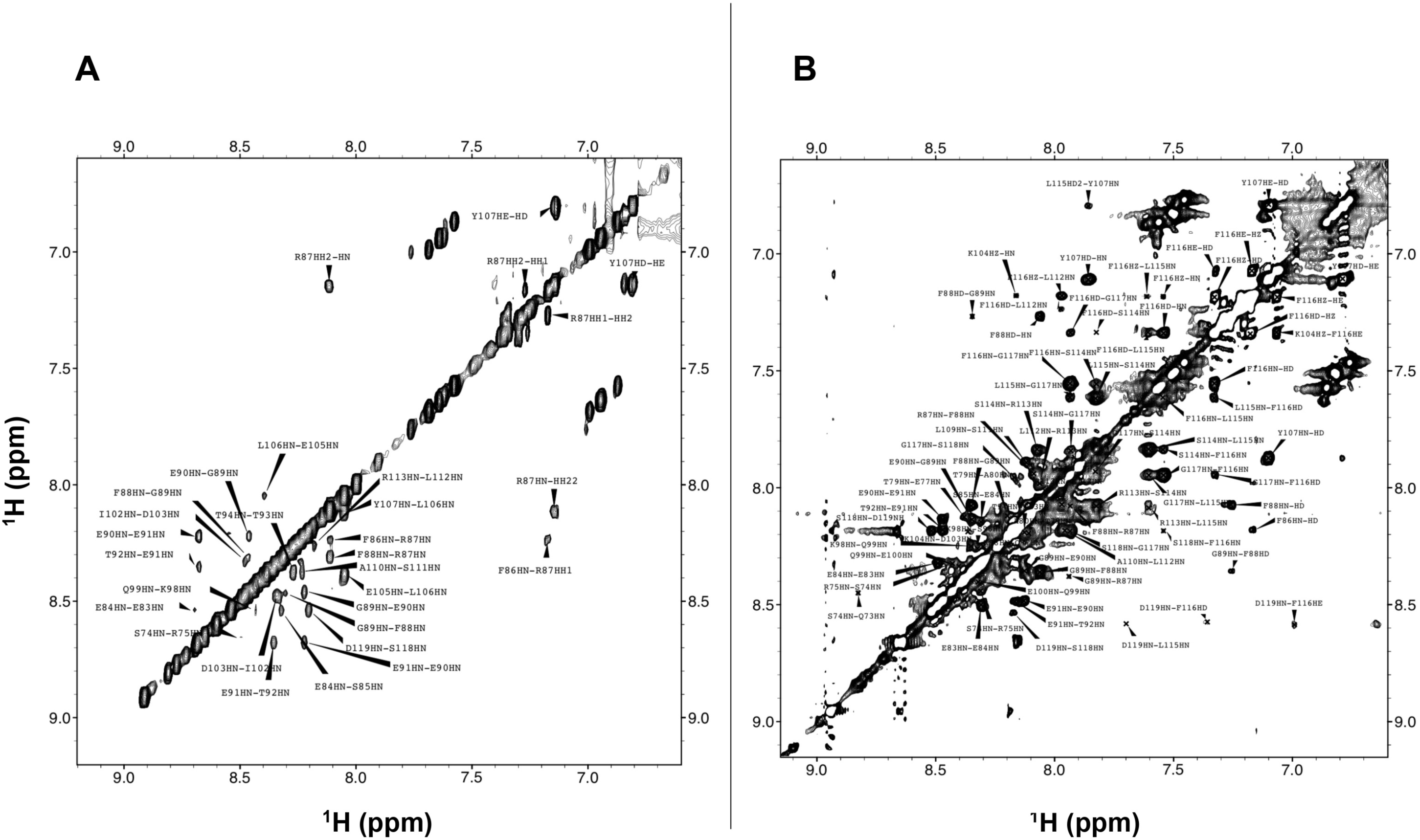
Proton NH and aromatic regions of 2D NOESY spectra of p6 recorded with a 150ms mixing time **(A)** at 10°C and in H_2_O, and **(B)** at 30°C with 450 eq. DPC-*d_38_* added. Most of the extra inter-residue NH-NH cross-peaks observed in the presence of DPC-*d_38_* are assigned to C-terminal residues.

**Figure 4.**
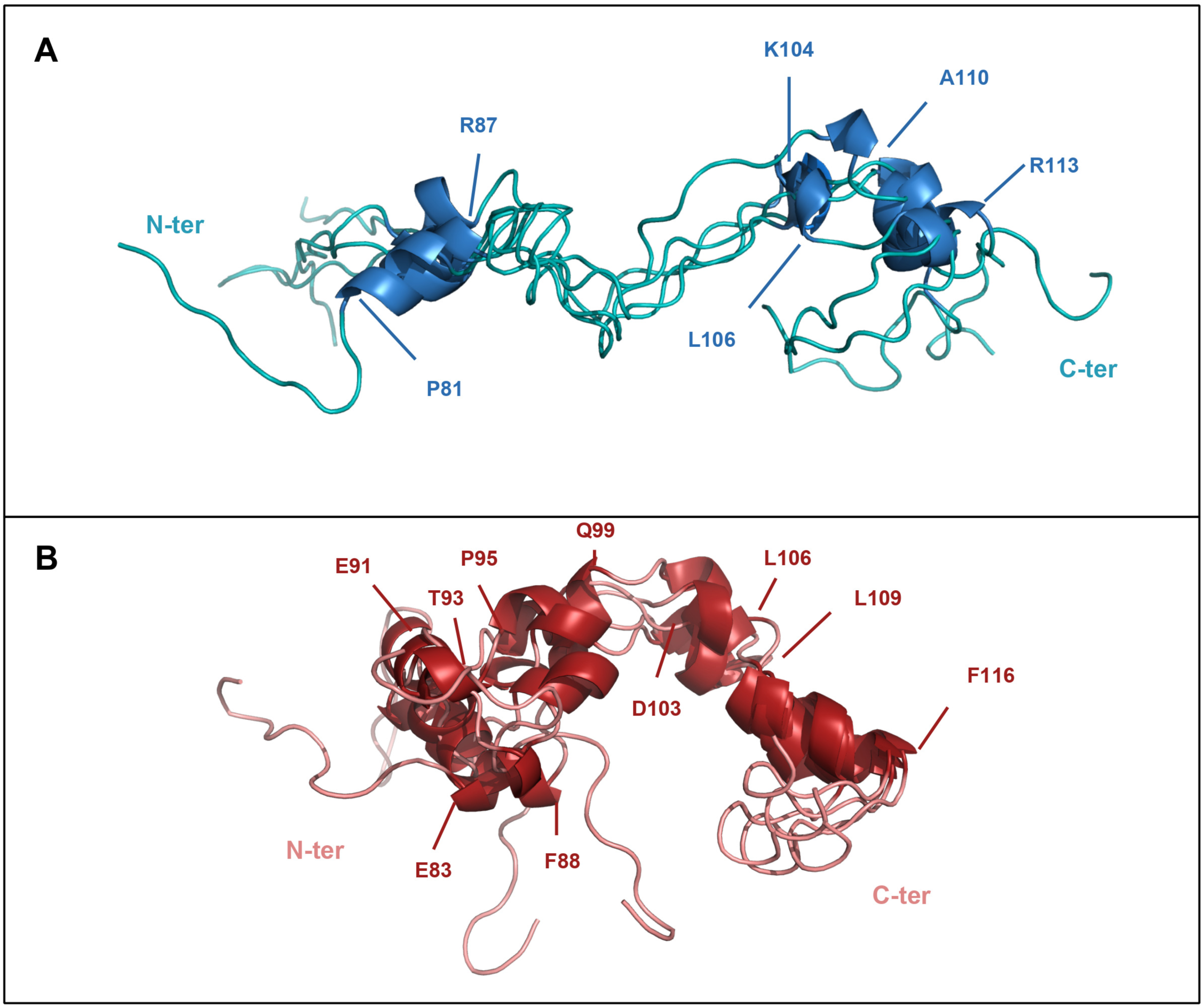
Backbone superposition of the solution structures of p6 computed with NMR restraints measured at 10°C and in H_2_O (**A,** in teal) or at 30°C and in 450 eq. DPC-*d_38_* (**B**, in salmon). In water p6 is in a flexible extended conformation with three small helices (in sky blue: P81-R87; K104-L106; A110-R113). In 450 eq. DPC-*d_38_*, p6 is more compact and folds around four to five helices (in firebricks : E83-F88; E91-Y93; P95-Q99 ; D103-L106 ; L109-F116).

While in water and at 10°C all p6 prolines share a *cis* conformation, the 2D NOESY spectrum recorded in DPC-*d_38_* and at 30°C exhibits correlations typical of a *trans*-conformation for proline P108 (HN-P107/HD1-P108 and HN-P107/HD2-P108; Fig. S5B). Such influence of temperature and solvent on the relative proportions of *cis*- and *trans*-isomers has been observed in the past on HIV p6 (57).

To get a deeper insight in p6 structure and dynamics, we also performed NMR experiments in DPC-*d_38_* on a sample of p6 partially ^15^N-labeled on the 6 amino acids F86, F88, S111-L112, L115-F116. For simplicity, this sample is called p6**^L^** hereafter, where L stands for labeled. Because they are clustered into two groups (N-ter side and C-ter side) that are far away in the p6 sequence, we have decided that they could provide interesting probes for the present studies. The [^1^H-^15^N] HSQC spectra of p6 **^L^** display strong chemical shift perturbations upon addition of 450 eq. DPC-*d_38_*, for S111, L112, L115 and F116, *ie* in the C-terminus (Fig. S5A, red and green cross-peaks). The CSPs are much weaker for F86-F88. The same experiments conducted in natural abundance at 10°C with an unlabeled sample of p6 (Fig. S4C) confirm the stronger variations observed for the C-ter part. S114-L115-F116-G117-S118-D119 display medium to strong CSPs. Medium CSPs are observed for G89-E90, K98-E100 and K104-Y107. At 10°C and in DPC, 10 peaks of correlation disappear from the [^1^H-^15^N] HSQC recorded in natural abundance, of which those of I102, E105, L109, A110 and R113. These last peaks could be recovered by raising the temperature to 30°C. We thus hypothesized that the origin of their line-broadening to baseline is more probably the consequence of interactions with DPC than to a transient loss of structure of the corresponding region. In the light of these results, it appears that the addition of DPC at 30°C has stabilizing effects on the weakly structured modules of p6.

### The ^15^N-labelled NC(p7) / p6 mixture

Protein p6 was gradually added to NC(p7)_1-55_either in water or in DPC buffer, and the perturbations of the ^15^N-H chemical shifts were followed on the [^1^H-^15^N] HSQC spectra. The CSPs induced by the addition to 10 molar eq. of p6 in water are displayed on Fig. S6. In water, the poor peak resolution of K11, V13, R32, K34 does not allow reliable CSP measurements. Interestingly, the correlation peak of Q2 appears upon addition of 1 equivalent of p6 (peak at 8.92 ppm ^1^H and 124.00 ppm ^15^N for 2.5 eq. p6; Fig. 5A). Its neighbor K3 undergoes one of the strongest CSP in the presence of 10 molar equivalents p6. In the N-terminal helix, G4, F6 and R7 undergo significant CSPs. Residues F16, N17, A25 and K26 in the one hand, and G40, H44, M46, K47, D48 and C49 in the other, located respectively in the zinc fingers ZF1 and ZF2 are also sensitive to the presence of p6.

**Figure 5.**
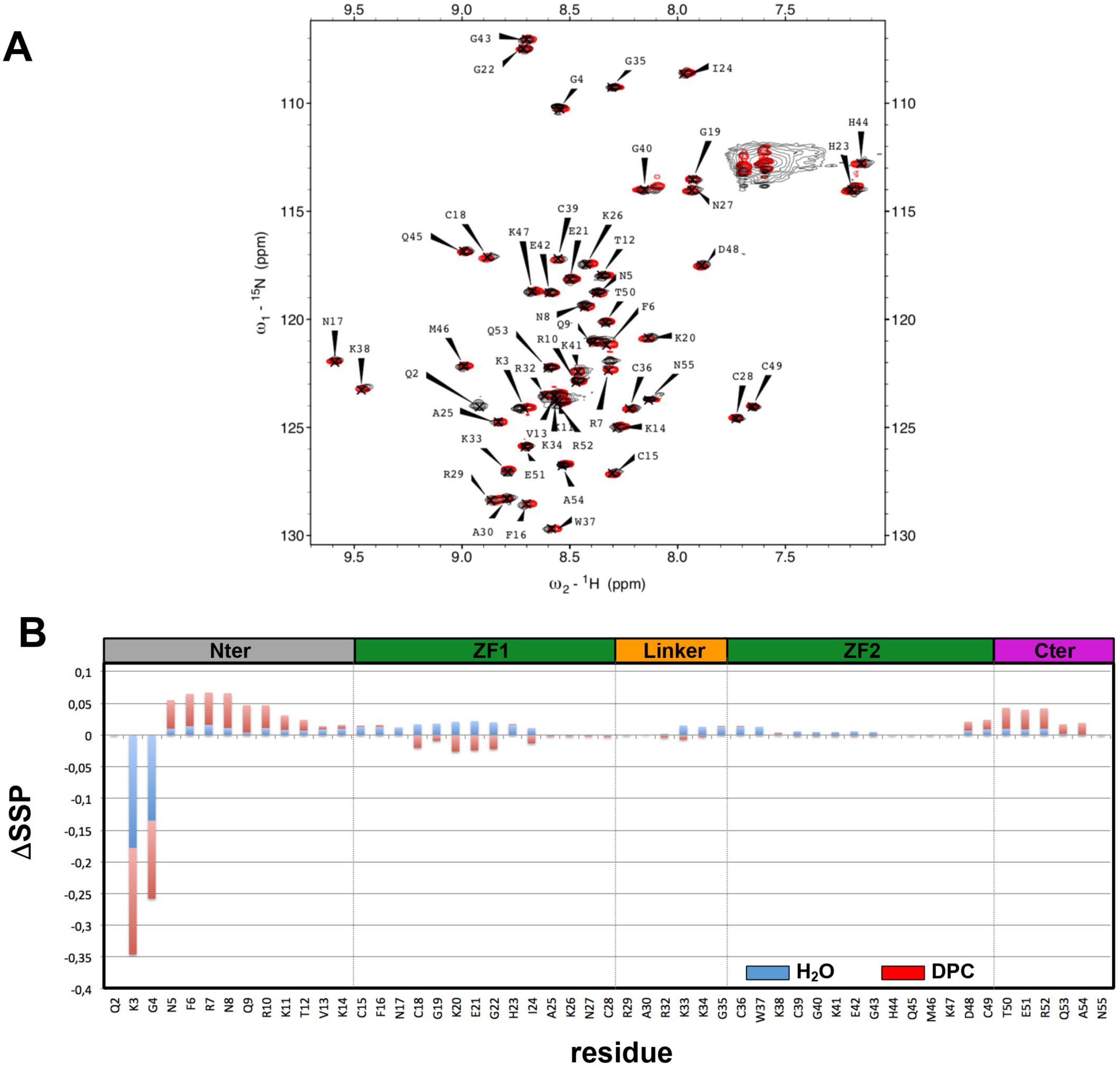
Influence of the addition of p6 on NC(p7)_1-55_. **(A)** Superposition of [^1^H-^15^N] SOFAST-HMQC of ^15^N-NC(p7)_1-55_ spectra recorded at 10°C as a free protein in H_2_O (red) and with 450 eq. DPC*-d_38_* in the presence of 2.5 eq. p6 (black). Q2 correlation peak is observed only in the complex. **(B)** Differences of NC(p7)_1-55_ Secondary Structure Propensities (SSPs) for the 1:10 complex with p6. (**Blue)** ΔSSPs measured in H_2_O at 10°C, using as reference the free state. (**Red)** ΔSSP measured in 450 eq. DPC-*d_38_* at 10°C, with reference taken in the presence of 0.5 eq. p6.

As noticed before, in the presence of DPC micelles and at 30°C, many amide cross-peaks (K3-G4-N5-F6-R7-N8-Q9) of the free ^15^N-NC(p7) _1-55_ are missing on the ^1^H-^15^N HSQC spectra (Fig S1B). Interestingly, most of these missing resonances reappear in the presence of 0.5 eq. of p6 added. Consequently, to monitor the effect of p6 on the whole structure of NC(p7)_1-55_, we have decided to compute the CSPs taking as reference the chemical shifts measured for 0.5 eq p6 (Fig. S6B), in addition to those measured without p6 (Fig. S6C). The N- and C-termini cluster most of the highest CSPs. The first one involves residues R10 to K14 located between the end of the N-terminal helix and ZF1, the second one T50, R52, Q53 and N55. The chemical shifts of K20 in ZF, K38 in ZF2 and R32 and K34 in the linker are also perturbated. These results show that the presence of p6 affects the entire length of NC(p7)_1-55_, with the highest effects observed on the two flexible termini.

We then compare the variation of the SSPs per amino acid induced by the presence of p6, in water and in DPC micelles. In DPC-*d_38_*, like for CSPs, we used the 0.5 eq p6 condition as reference (Fig. 5B). The addition of p6 triggers stronger effects in the presence of DPC-*d_38_*, particularly at the N-terminal region, leaving both the linker and ZF2 mainly unaffected. Conversely, opposite destabilizing effects are observed in DPC-*d_38_* in the region ranging from C18 to I24 which corresponds to the first zinc finger.

### The p6^L^ / NC(p7)_1-55_ mixture

We have monitored the effect of the addition of NC(p7)_1-55_ on p6^L^ in 450 eq. DPC and at 10°C. The labeled residues, F86 and F88 on the one hand, S111, L112, L115 and F116 on the other are clustered into two groups (N-ter side and C-ter side) that are far away in the primary sequence. They provide interesting probes to test the interaction of p6 termini with NC(p7)_1-55_. The [^1^H-^15^N] HSQC spectra of free p6^L^ recorded in water (red cross-peaks) and in DPC-*d_38_* (green cross-peaks) with the spectrum recorded in the presence of both DPC-*d_38_* and 1 eq. NC(p7)_1-55_ (black cross-peaks) are displayed in Fig. S5. Significant shifts are visible for S111, L115 and S116 (shifts from green to black positions) on the contrary of F86 and F88. They are however much smaller than those induced by the addition of DPC-*d_38_* (shifts from red to green positions) on the free protein.

### Quantification of the interaction between NC(p7)_1-55_ and p6 in aqueous and micellar conditions

Chemical shifts perturbations were observed on [^1^H-^15^N] SOFAST-HMQC spectra upon the gradual addition of p6 to ^15^N-NC(p7)_1-55_. They are more important in DPC than in water (Fig. S6). To obtain a preliminary estimate of the interaction strength between the two partners in DPC, we monitored the evolution of CSPs as a function of p6 concentration. We restricted our analysis to correlation peaks that met the following criteria: visible in the absence of p6, good resolution across all the titration, CSPs above the average. We also choose not to consider the C-terminal N55. This reduced the list to the peaks of 10 amino acids (Fig. 6A): F6, R7 and R10 for the N-terminal region, K14, N17 and I24 from ZF1, A30 for the linker, K38 from ZF2 and for the C-terminal E51 and R52. The experimental data recorded at 10°C (not shown) were fitted according to Equation 2 (see materials and methods). They are consistent with apparent dissociation constants (K_dapp_) in the order of few hundreds of micromolar: 119 μM (F6), 81 μM (R7), 28 μM (R10), 356 μM (K14), 342 μM (N17), 468 μM (I24), 1.76 mM (A30), 454μM (K38), 803 μM (E51) and 391 μM (R52). The measured K_dapp_ vary depending on the amino acid, and reflect the binding of p6 as well as any changes in local conformation. The smaller computed K_dapp_ are for F6, R7 and R10, which lie in the flexible α-helix region strengthened by the presence of p6. At the opposite, the largest K_dapp_ are observed in the flexible regions of the protein, at the C-ter part and for the linker residue A30.

**Figure 6.**
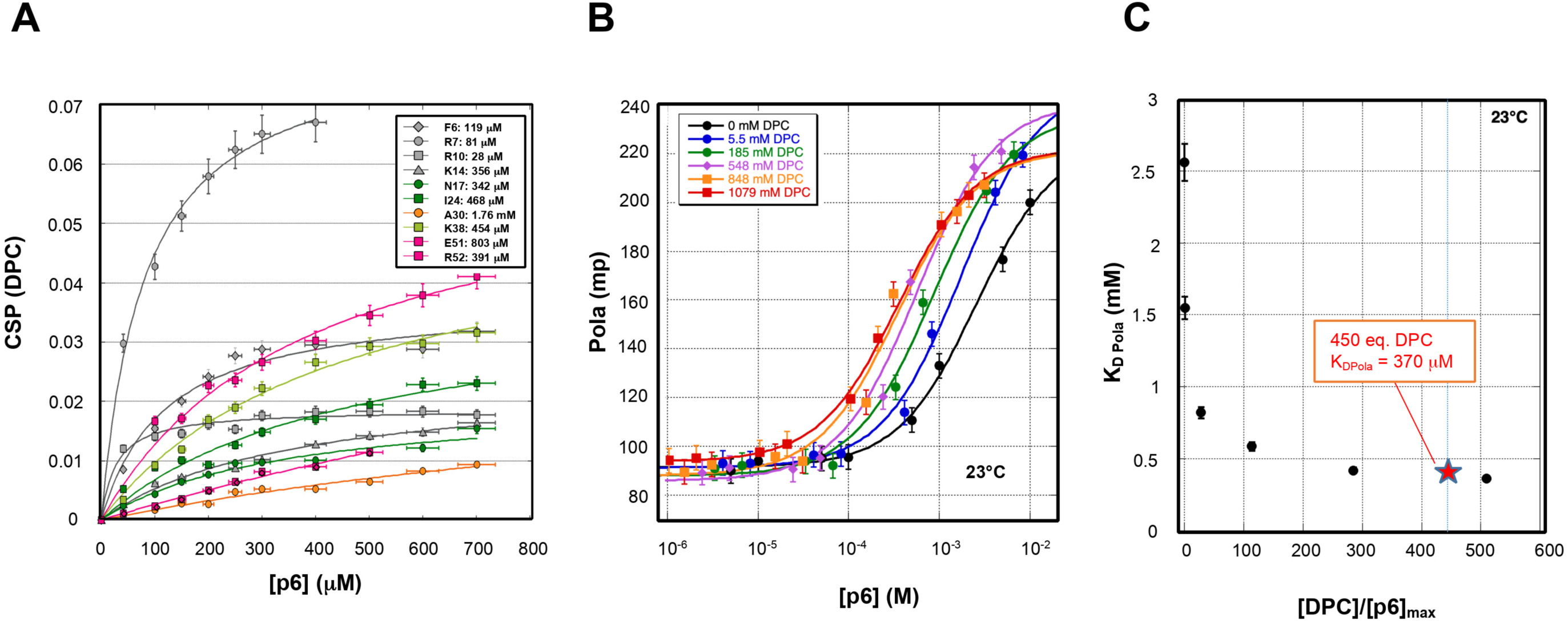
Interaction between NC(p7)_1-55_ and p6. The experimental data were fitted according to Eq. 2. **(A)** NMR K_d_ values computed from the variation of CSP upon addition of p6 and observed on [^1^H-^15^N] SOFAST-HMQC experiments for 10 amino acids (F6, R7, R10, K14, N17, I24, A30, K38, E51, R52). Error bars on CSPs were set to 5% and on p6 concentrations to 1%. (**B,C)** Fluorescence polarization recorded at 23°C on NC(p7)_1-55_ as function of p6 and DPC concentrations. Error bars were set to 5%. **(B)** Superposition of the different titration curves recorded for DPC concentration ranging from 0 to 1.08 M. **(C)** Evolution of K_d_ as a function [DPC]/[p6]_max_.

Fluorescence anisotropy enabled us to monitor the titration over a broader range of p6 concentrations. As before, the titration of NC(p7)_1-55_ by p6 was monitored in both NMR buffers, with and without the addition of DPC. Two sets of experiments at 23°C were conducted using two different NC(p7)_1-55_ concentrations (1.3 and 10 μM), by recording the NC(p7)_1-55_ tryptophan fluorescence. Fluorescence intensities were insensitive to the addition of p6 without or in the presence of DPC (not shown), which is consistent with the absence of noticeable variations of CSPs and SSPs for W37. Titrations were thus followed by recording perpendicular and parallel intensities from which polarizations were computed, for concentrations of p6 ranging from 0 up to 1000 eq. NC(p7)_1-55_ and for increasing concentrations of DPC from 0 to 1.079 M (Fig. 6B). In the absence of DPC, data analyses according a 2-step model yield a K_d_ value between 2.43 and 2.66 mM with best fit obtained for 2.56 mM. In Equation 2, [p6]_max_ was set to the maximum p6 concentration reached in the titration. Limited accuracy in determining K_d_ stems largely from the inability to reach high enough p6 concentrations to fully saturate the binding during titration. Nevertheless, data clearly show that p6 slightly interacts with NC(p7)_1-55_ even in the absence of any DPC. In Fig. 6C, the DPC concentrations were varied and expressed as molar equivalents of [p6]_max_. We observe that the computed K_d_ strongly decreases up to a DPC concentration of 100 eq. of [p6]_max_ to reach a plateau above 200 eq. of [p6]_max_, corresponding to a lower K_d_ value of about 370 mM. This value is close to the value estimated from the CSP titrations. The fluorescence anisotropy data show that the affinity of p6 for NC(p7)_1-55_ increases in the presence of lipids and that the K_d_ decreases by a factor of about 7 in the presence of DPC micelles. Hydrophobic surrounding thus strengthens the interaction between NC(p7)_1-55_ and p6, which albeit weak, preexists in water.

### Modeling of the interaction between NC(p7)_1-55_ and p6

*In vivo*, p6 and NC(p7)_1-55_ domains are connected by the flexible SP2. To determine whether the interaction between the two free domains detected by NMR could be transposed to an intramolecular interaction in the context of Gag or in the mature NC(p15) form, we refined a NC(p15) 3D model generated by AlphaFold3 using Xplor-NIH software and EEFX force field. The same set of restraints as described before was employed, including NMR-derived distance restraints for both p6 and NC(p7)_1-55_. As the largest DSSPs in the mixture were observed in the N-ter residues of NC(p7)_1-55_ and in the C-ter residues of p6, we postulated that these regions form an interaction interface in the 1:1 complex. Therefore, artificial distance restraints (4–25 Å, at residues 6-11 for NC(p7)_1-55_ and 111-116 for p6) were applied between the two helices during the refinement procedure to favor an antiparallel arrangement. Two hundred structural models of the intramolecular complex were generated using the experimental restraints, corresponding to either water or 450 eq DPC*-d38* buffer conditions.

Superpositions of the 10 lowest-energy models consistent with the experimental restraints are shown in Fig. 7. In both cases, the 3D models confirm that one p6 can fold back and cover the entire length of NC(p7)_1-55_, with an interaction between its C-terminal pseudo-helix formed by residues F116-L115-S114-R113-L112-S111 and residues F6-R7-N8-Q9-R10 of the NC(p7)_1-55_ N-terminal helix. In all models, NC(p7)_1-55_ adopts an extended conformation, with F16 and W37 unstacked and therefore in the open state (data not shown), in contrast to the initial AlphaFold3 structure, where the two aromatic side chains are found in the closed conformation. In the intramolecular complex, p6 interacts with different faces of the N-terminal helix of NC(p7)_1-55_. In DPC*_-d38_* buffer, its localization appears more clustered around the N-terminal helix than in water, consistently with the lower Kd measured under hydrophobic conditions.

**Figure 7.**
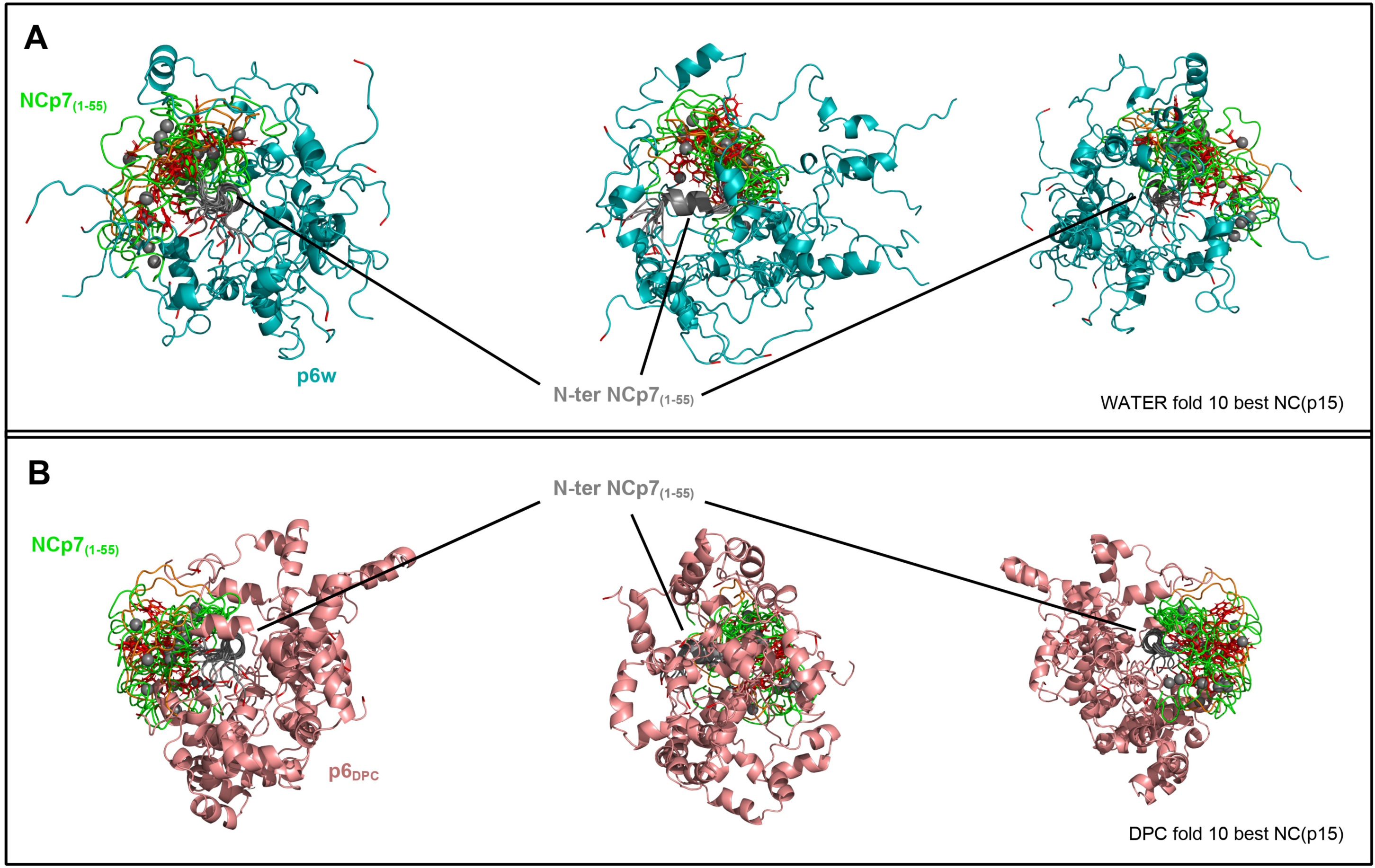
Superposition of the ten lowest-energy models of NC(p15) refined with NMR restraints in water (A) or DPC micelles (B), showing intramolecular interactions between the NC(p7)_1-55_ and p6 from different views. All models were superimposed onto residues 5-10 of the N-terminal domain of NC(p7)_1-55_ (gray). NC(p7)_1-55_ zinc fingers are shown in green, and F16 ad W37 in red. For clarity, the NC(p7) C-terminal TERQAN residues and the adjacent SP2 linker are omitted. (A) Intramolecular complex in water with p6 (in cyan). (B) Intramolecular complex in DPC with p6 (in salmon).

## DISCUSSION

The nucleocapsid NC(p7)_1-55_ protein is a small protein characterized by two CCHC-type zinc fingers connected by a flexible linker. The N- and C-termini and the linker are rich in basic residues, enabling the protein to interact with various nucleic acids. Molecular dynamics simulations and Density Functional Theory (DFT) studies have shown that NC(p7) interacts with RNA and DNA primarily through electrostatic interactions, facilitated by its basic residues (58). NC(p7) plays multiple key roles in the Human Immunodeficiency Virus 1 (HIV-1) life cycle, orchestrating various processes such as reverse transcription, RNA dimerization, and genome packaging. It chaperones the RNA reverse transcription into double-stranded DNA (dsDNA). It also selects, condenses and protects the viral gRNA for packaging. Understanding NC(p7)_1-55_ structural and dynamic properties is crucial for elucidating its different roles in the viral HIV cycle. p6 is a small acidic protein. As part of Gag, it lies in the vicinity of NC(p7) separated only by the flexible SP2 linker. This spatial proximity led us to question whether the two components could interact, and if so, how and with what strength. We have thus studied the structures and the dynamics of NC(p7)_1-55_ and p6 as free proteins and as in complex, in acetate aqueous buffer, both in the presence or the absence of DPC.

### NC(p7)_1-55_ adopts a dynamic compact conformation, with a small helix at its flexible N-terminus

The structure of NC(p7) is known to be partly folded around two zinc fingers (ZF1 and ZF2) which are separated by a flexible linker (39, 46). The N-terminus is very basic and adopts a helix-like structure in the presence of RNA (44, 45) which has never been shown on the free protein in solution. In the first part of this manuscript, we evidenced, in aqueous conditions at pH 6.5, the formation of a small α-helix within the flexible the N-terminus of free NC(p7)_1-55_, ranging from N5 to R10.

NMR models of structure were computed with distances and angles restraints (Fig. 2B). Like observed before with the shorter NC(p7)_13-55_ (39, 46), our 3D models of structure all exhibit compact conformations organized around the two CCHC zinc fingers which are close in space. This is experimentally supported by the observation of long-range NOEs on the [^1^H–^15^N] 3D NOESY-HSQC experiment recorded with a mixing time of 150ms, between the side chains of residues that are distant in sequence: Phe16/Lys38, Asn17/Lys38, Trp37/Cys18, Trp37/Phe16. Spin diffusion alone could not account for these NOEs. The two aromatics residues F16 and W37, despite not stacked like in the structure of Morellet *et al.*, adopt a “closed” conformations like described in the literature. The dynamic behavior of the linker and the two zinc fingers has been described in many studies (53). It is now commonly admitted that open and closed conformations coexist because of the linker dynamics. Their difference of free energies are very small, computed to be less than 0.75–1.9 kcal/mol depending on the protein sequence and the ionic strength (59). The force field we used for folding and refinement during protein structure modeling (EEFX) harbor differences in the energy terms, especially for the non-bonded contributions compared to those used by Morellet *et al.* (39). It is thus not surprising that the computed structures present local differences in the protein regions that are quite flexible and highly charged, which is the case of the two flexible linkers, RKTVK between the α-helix and ZF1 (pI≈11) and RAPRKKG (pI≈12) between ZF1 and ZF2. These two linkers are amongst the most positively charged regions of NC(p7)_1-55_. These differences have repercussions on the relative positions of the helix and ZF1 in the one hand and of ZF1 and ZF2 in the other.

Our models of structure all disclose a small α-helix of about one turn, ranging from F6 to R10 with a helical pitch of 5.4 Å. The orientation of the N-terminus and thus of the helix with respect to the zinc fingers is variable. Recent NMR relaxation techniques experiments (CPMG and CEST) reveal that the NC domain of Gag exists in minor conformers across all its maturation forms (NC(p15), NC(p9), and NC(p7)). These minor conformers involve only ZF2 rearrangements. However, large Rex values were observed in the flexible N- and C-terminal regions of NC(p7) at 35°C and pH 5.5, suggesting a fast exchange of amide proton with solvant and a presence of temperature-dependent conformational dynamics in these regions (60). Thus, to our knowledge, our NMR data bring experimental evidence of a small α-helix for the free NC(p7)_1-55_ domain, in the absence of any nucleic acid partner.

We tested the effect of a temperature increase on ^15^N-NC(p7)_1-55_ by recording [^1^H–^15^N] TROSY experiments at 10°C and 30°C (Fig. S1A). At 30°C and in water, 12 correlation peaks disappear from the spectrum (53 NH peaks observed at 10°C *vs* 41 at 30°C) confirming the poor structural stability of the corresponding regions and particularly of the N-ter region which displays the disappearance of eight peaks (K3 to R10). The other peaks are assigned to R29, R32, K34 (inter finger linker) and G43 and Q53 (C-ter part). This shows that these regions rapidly exhibit changes of conformational exchange regime and that they are weakly observed consistently with enhanced T1 and T2 and decreased hetero-NOEs for R32 and at the C-terminus (Fig. S3).

The influence of the solvent hydrophobicity on the dynamics of the protein was investigated by measurements of the T1 and T2 relaxation times in the presence of 450 eq. DPC-*d_38_* and of the [^1^H–^15^N] hetero-NOE values (Fig. S3). The differences observed along the sequence starting from ZF1 up to the C-terminus are not statistically significant. Importantly, like in water, the results do not highlight any differences of stability between the two zinc fingers. Unfortunately, the [^1^H–^15^N] HSQC correlation peaks of the residues the N-terminal α-helix are not seen in DPC-*d_38_* (Fig. 2A) with the consequence that neither the structure nor the dynamics of this region in the free protein could be probed. Nevertheless, this observation suggests a potential interaction of the N-terminal of part NC(p7)_1-55_ with the DPC micelles. The observation that the T1 and T2 relaxation times of R10 are lowered in the presence of DPC-*d_38_* and fall within the range of the values observed for the central part of the protein, suggests an increase in rigidity or a reduction of the possible conformations for the linker between the α-helix and ZF1.

### p6 folding is enhanced in the presence of DPC, particularly at its C-terminus

The structure of free p6 in solution was computed under NMR restraints measured in water and in DPC-*d_38_* (Fig. 4). The protein is highly dynamic with small helices. Their occurrence raises in the presence of DPC and is favored by an increase of temperature. They are close to TALOS predictions (Fig. S4). Notably, both computations disclose one α helix at the C-terminus which is a little bit more extended in the presence of DPC micelles, consistently with the higher number of corresponding inter-residue NH-NH cross-peaks observed on the NOESY spectra (Fig. 3). Solbak *et al.* previously demonstrated that p6 adopts a structured conformation with N- and C-terminal helical domains for 100 mM DPC at pH 7, without the use of organic co-solvents (50). Using surface plasmon resonance (SPR) spectroscopy, the authors show that p6 directly interacts with a cytoplasmic membrane, involving both its N- and C-terminal regions. Like mentioned by Solbak *et al.*, we observe that the p6 helical content is increased by the presence of DPC micelles. We observed two extra helices spanning residues E91-T93 and S96-Q99, which do not preexist in the aqueous solution structure in the absence of DPC (Fig. S4D). The enhanced helicity observed in our study may result from the higher DPC-to-protein ratio we employed (450 eq.) compared to the 50-100 equivalents used by Solbak *et al.*.

### NC(p7)_1-55_ interacts with p6 and the interaction is stronger in the presence of DPC micelles

We provide the first evidence of an interaction between the two proteins in solution, and particularly in lipidic conditions, using NMR and fluorescence anisotropy. The interaction is weak in water (K_d_ about 2.5 mM at 23°C) but real and supported by concentration-dependent changes in the chemical shifts of the amide NH correlation peaks in [^1^H–^15^N] SOFAST-HMQC experiments (Fig. 6). The ΔSSPs measured on NC(p7)_1-55_ in water and induced by the presence of p6 are weak and distributed across almost the entire sequence of the protein. However, the appearance of a correlation for Q2 in the [^1^H–^15^N] HSQC spectrum as soon as 1 equivalent of p6 is added, discloses a change of exchange regime, consistent with an interaction between the N-terminus and p6 (Fig. 5).

The affinity appears to increase with higher DPC concentrations, to reach a K_d_ plateau of 370 μM above the CMC at 23°C (Fig. 6C) or measured K_dapp_ close to 250μM by NMR at 10 °C (Fig 6A). This is also supported by the observation that the largest NC(p7)_1-55_ ΔSSPs mainly affect the terminal residues and in a slighter extent those in the ZF1 region (Fig. 5B). The signs show that the presence of p6 in DPC stabilizes the N-ter and the C-ter parts of NC(p7)_1-55_ while destabilizing ZF1 and part of the neighboring linker. The C-terminus of the free NC(p7)_1-55_ is unstructured. We show that the presence of p6 restricts its movements leading to the observation of positive ΔSSPs. Because the strongest CSPs observed for p6 upon addition of NC(p7)_1-55_ are clustered around its C-ter part (Fig. S4A), it reasonable to assume that the C-terminus of p6 interacts and stabilizes the NC(p7)_1-55_ N-terminal α-helix. Altogether, our results highlight the adaptation of the NC(p7) protein structure to solvent hydrophobicity and protein partners.

### 3D models of the NC(p7)_1-55_ : p6 interaction

To get insight on how NC(p7)_1-55_ and p6 interact at the molecular level, we have run simulated annealing and refinement protocols using as restraints the experimental distances measured on the free partners. The simulations of the intramolecular complex in water and in DPC differ primarily in the experimental restraint sets applied to p6. To get closer to the natural context of Gag, we started from a NC(p15) 3D model generated by AlfaFold3, in which NC(p7) is correctly structured around its two zinc fingers and p6 partially structured around small helices.The refinement temperature was limited to 1500 K to preserve the overall structural features while allowing limited conformational changes driven by the experimental restraints. Because no experimental intermolecular NOEs were observed between the two partners (most probably as a consequence of a relatively high Kd), we implemented the experimental restraints by a set of artificial intermolecular distances ranging from 5 to 20 Å between the N-terminal helix of NC(p7)_1-55_ and the C-ter part of p6. These artificial distance restraints were easily accommodated in the 3D models of lower energy computed with Xplor-NIH/EEFx (Fig. 7). In the computed 3D models, the C-terminus of p6 gets close to the N-ter α-helix of NC(p7)_1-55_ with different helix-helix orientations (Fig. 7A,B). In the cell, such intermolecular interaction may be driven by the opposite charge of these two regions, highly basic for NC(p7)_1-55_, acid for p6. In both simulations, the p6 C-ter part interacts with NC(p7)_1-55_ from the same side relative to the two zinc fingers position (Fig. S7). However, the results suggest that the conformational space sampled by p6 is broader in the absence than in the presence of DPC micelles. Together with the lower K_d_ measured in DPC and the higher resonance shifts observed, this suggests a mechanism in which, as p6 gets close to the membrane, its back-folding onto NC(p7)_1-55_ is favored and its motions around the N-ter helix become more restricted.

### p6, a competitor of gRNA for the binding to the nucleocapsid ?

The NC(p7) structure exhibits conformational flexibility both as a free protein and in complex. The positively charged regions including the N-terminus and the inter-ZF linker, can establish electrostatic contacts with polyanions like nucleic acids. The two zinc fingers, containing the hydrophobic residues F16 and W37, create a stacking platform that preferentially interacts with the unpaired bases of nucleic acids, especially with guanines. However, they display distinct behaviors depending on whether the interacting nucleic acid is DNA or RNA. During the initial phase of the interaction with the viral DNA, a guanine is inserted within the hydrophobic zone of ZF2 by interactions with W37, thereby leaving ZF1 available to engage in subsequent interactions with a second G, T or C base (61). The RNA recognition mode is different, with ZF1 and ZF2 both interacting with two Gs separated by a third residue (G-X-G). Dannull *et al.* have showed that the N-terminal part of NC(p7), comprising the first zinc finger, is sufficient for RNA binding (58) (62). ZF1 is therefore essential for the recognition of viral RNA, in contrast to DNA recognition. Recent studies (63) show that the mutations K3A/F6A/Q9A mostly abrogate ssRNA aggregation but maintain DNA condensation, suggesting that the N-terminus is also primarily involved in stabilizing the interaction network between the nucleocapsid and nucleic acids. Mutation assays demonstrate that K3, R7, R10, R29 and R32 but not K11, K14, K33 and K34, are essential for specific binding to RNA(62). ZF1 is similarly engaged in the context of NCp(15) and NC(p9), which both comprise two palindromic sequences in the N-terminus and in the SP2 spacer, [(K(3/59)-G(4/58)-x-F(6/56)-x-x-Q(9/53)-R(10/52)] that act as recognition elements for viral RNA binding. They are highly conserved amongst all HIV subtypes except at position 3 and are thought to establish bridges between two gRNA strands (63).

Because our NMR data show that the N-terminus and the ZF1 regions of NC(p7)_1-55_ are involved in the complex with p6, but not ZF2, we hypothesize that p6 can compete with the viral RNA and not DNA for binding to NC(p7), through its Psi packaging signal, which is composed of highly structured secondary stem-loop elements (SL1–SL4) essential for selective recognition and encapsidation by Gag. The structure of the complexes with the RNA hairpins SL2 (pdb: 1f6u) (44) and SL3 (pdb: 1alt) (45) have been published. In the two cases, the NC(p7) harbors a small N-terminal α-helix, and the zinc fingers interact with the apical loop of the partner RNA. This interaction brings the helix close to the RNA. To compare the relative positions of the RNA hairpins and of p6 and to take into account the fact that the conformational landscape of the N-terminus of NC(p7 or 15) in the Gag context is expected to be a little more restrained than in the free state, we have superimposed the residues 5 to 10 of the N terminal α-helix of NC(p7)_1-55_ in complex with SL2 (1f6u.pdb; (44)) and SL3 (1alt.pdb; (45)), and of our models of NC(p15) in complex with p6 (Fig. 8). The two RNA hairpins occupy distinct regions around NC(p7)_1-55_, particularly with respect to its N-terminal portion (Fig 8A). In water, p6 samples a broad conformational space that overlaps the regions occupied by SL2 and SL3, although localizations on the SL2 side appear to be slightly over-represented (Fig. 8B). In contrast, in DPC, the conformational space sampled by p6 is more restricted and largely confined to the region occupied by SL2 (Fig 8C). This suggests that the SL2 side would be favored, as p6 helical content increases consistently with an increase in the hydrophobicity of the environment. This also supports the assumption that p6 could compete the viral RNA for binding in the cell. It could destabilize ZF1, thereby hindering its association with non-essential RNAs while promoting the stacking of viral RNAs (interaction F16/W37 with GXG unpaired bases), which are known to exhibit higher affinity, with dissociation constants (K_d_) in the nanomolar range.

**Figure 8.**
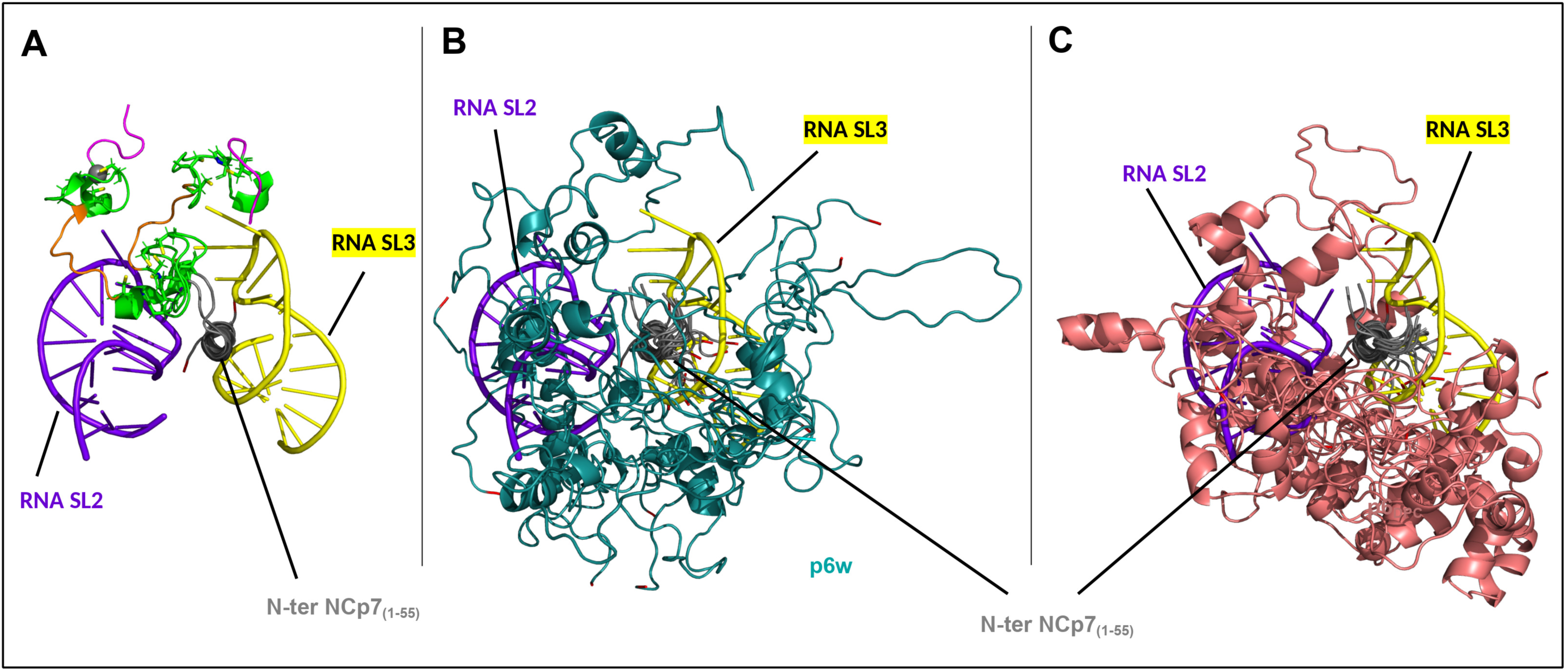
**(A)** Superposition, onto residues 5-10 of the N-terminal domain of NC(p7)_1-55_ (gray), of the NC(p7):SL2 (in purple, pdb: 1f6u) and NC(p7):SL3 (in yellow, pdb: 1a1t) complexes. **(B)** Lowest-energy models of NC(p15) refined with NMR restraints in water (10 models) have been superimposed onto NC(p7):SL2 (in purple, pdb: 1f6u) and NC(p7):SL3 (in yellow, pdb: 1a1t) complexes **(C)** Lowest-energy models of NC(p15) refined with NMR restraints in DPC (10 models) have been superimposed onto NC(p7):SL2 (in purple, pdb: 1f6u) and NC(p7):SL3 (in yellow, pdb: 1a1t) complexes. **(B, C)** For clarity NC(p7) zinc fingers, TERQAN and SP2 residues were omitted. Same color coding as in Figure 7 was used for p6.

We hypothesize that the N-terminus of NC(p7) can accommodate the simultaneous binding of distinct binding partners (p6 and nucleic acids), on the condition they target different sides of its N-terminal α-helix. In such model (Fig 9), p6 could switch from one conformation to the other to fulfill its task, competing with and thus preventing the binding of a nucleic acid on the one hand, and making available the other side of the NC(p7) α-helix for another partner. This model may help reconcile previously conflicting results demonstrating that p6Gag is important for intact gRNA selectivity towards the other multiple RNAs, being at the same time dispensable for RNA Psi recognition (64).

**Figure 9.**
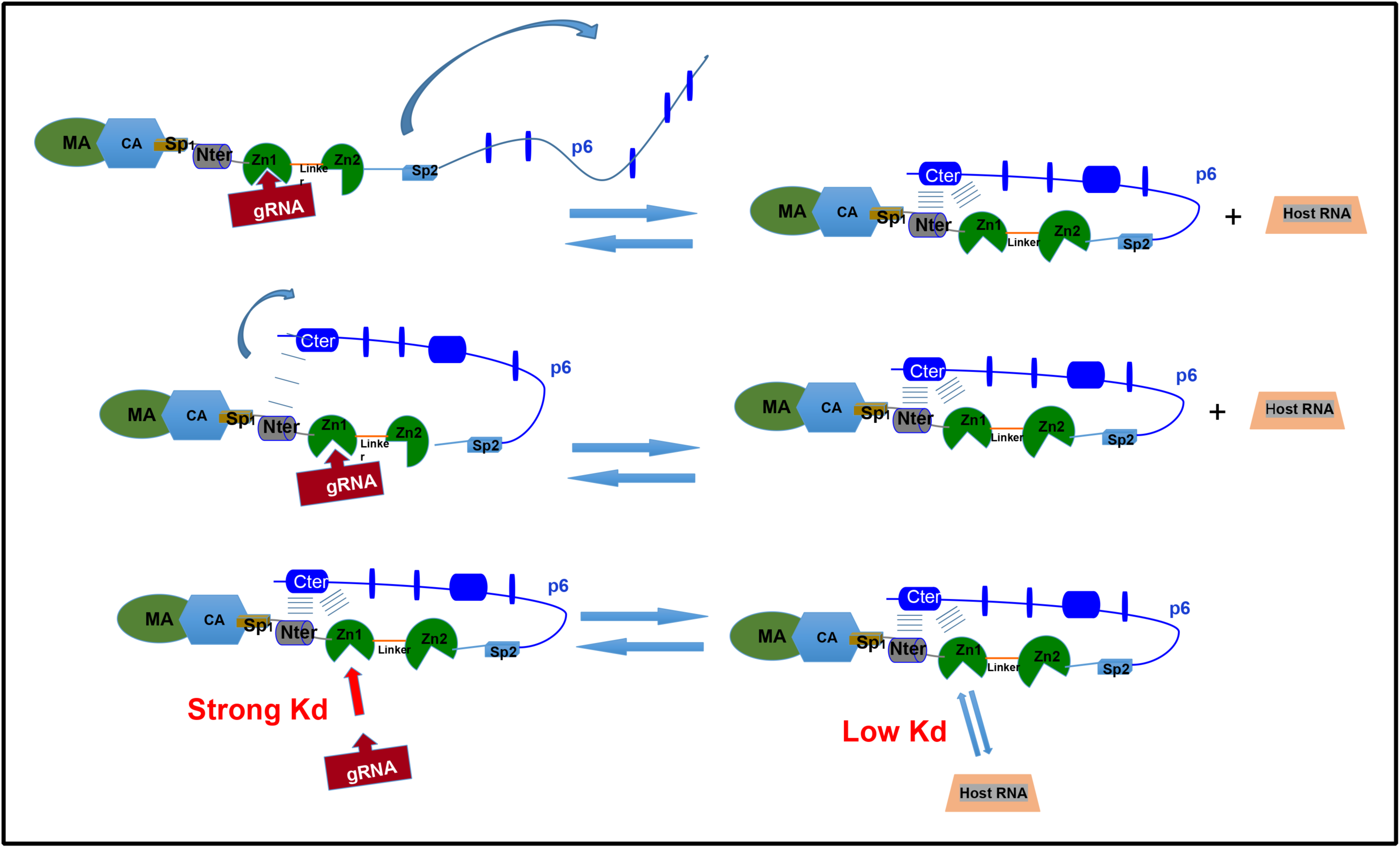
Schematic model based on our results. Our results indicate the possibility of an intermolecular interaction between NC(p7)_1-55_ and p6. In the context of full-length Gag, the occurrence of such an interaction is increased, as it corresponds to an intramolecular event between two regions that are in close proximity and connected by a flexible linker. The complex formed by the fold-back of p6 onto NC(p7) is expected to be dynamic. p6 would thus compete with all molecules for binding to NC(p7). During migration to the membrane, viral gRNA selection occurs via specific recognition by the first zinc finger. Our work shows that the interaction of p6 with NC(p7) induces its destabilization, which in turn may decrease its overall affinity for RNAs, particularly for non-cognate RNAs. The viral RNA, with reported dissociation constants in the nanomolar range, is therefore expected to efficiently compete with p6 for association with NC(p7) (intermolecular K_d_ in the order of few hundreds of micromolar) and disrupt the helix-helix interaction between the termini of these two proteins.

### The Ncp7-p6 in the cell

Our results suggest that in the cell or in the virion, the NC(p7)_1-55_ and p6 domains can interact. From *in vitro* acquired data and molecular simulations, we showed that p6 can compete with nucleic acids to bind NC(p7)_1-55_. In the infected cell and before the budding of the virion, NC(p7) _1-55_ is connected to MA-CA-SP1 on its N-terminus, and SP2-p6 on its C-terminus. All these proteins together form the Gag polyprotein. *In vivo*, NC(p7) and p6 are thus both domains of the same dynamic Gag polyprotein. They are connected by the flexible linker SP2, enabling them to adopt spatially proximal conformations during a fraction of the time.

In a recent study, large-scale all-atom molecular dynamics simulations were carried out on full-length Gag proteins in different multimerization states, bound to an asymmetric lipid membrane (65) and using a starting model reconstructed from cryo-ET and NMR structures. The inter-domain interactions of Gag were assessed using a heterogeneous elastic network model (hENM) (66). (65). The p6 domain is interestingly back-folded on it with various orientations and its C-ter helix close to either ZF1 or the N-terminus of NC(p7). Albeit the authors do not report any strong interaction between p6 and NC(p7), their models are reminiscent to those we obtained mixing our NMR results on the two separate domains and molecular modeling in the NC(p15) context (Fig. 7 and S7). In the cell, Gag thus most probably exists in an equilibrium of open and closed forms of the NCp(7)-SP2-p6 C-terminal domain. Almost no free p6 can be evidenced in the cell, most probably because the few small helices embedded in its intrinsically disordered structure are to weak to resist to the cellular proteases. The interaction with the NC(p7) domain, reinforced by its intramolecular nature in the Gag context, likely confers protection to the p6 domain against degradation. The Gag-gRNA complex starts to form in the cytoplasm and proceed at the membrane until completion of the virion assembly and budding (64). In water and in the absence of DPC, we measured weak affinities between NC(p7)_1-55_ and p6. This suggests that intramolecular interaction between p6 and NC(p7) within Gag, albeit weak, is possible and could occur, most probably, transiently and may be statistically significant enough to repel the unwanted nucleic acids, thereby ensuring that the NC(p7) region of part of the Gag population remains available to interact with the viral nucleic acids (Fig 9).

Gag is also known to recruit latter on two copies of single stranded and unspliced gRNA for the assembly of the future virion. Our results suggest that p6 can target the same NC(p7) regions than the gRNA SL2 and SL3 hairpins of the Psi packaging signal (Fig. 8), with an enhanced strength in the vicinity of the membrane. We can then assume that p6 may be an excellent competitor of the non-specific nucleic acids, but not strong enough to prevent the binding of the gRNA to Gag via NC(p7). The interaction between p6 and NC(p7) would thus indirectly participate to the selective recruitment of the gRNA (Fig. 9). Gag can form dimers and trimers but also hexamers that are assembled by their CA_CTD_-SP1 domains through a six-helix bundle (68) *In vitro*, a high affinity RNA oligomer can promote a Gag dimerization stronger than that through the CA_CTD_-SP1 domains, suggesting that the oligomeric state of Gag can be influenced by the presence of the gRNA (69). If p6 competes with the gRNA at this stage, this means that it may indirectly regulate Gag oligomerization.

Our study can also make sense in a step where both NC(p7) and p6 exist as domains part of smaller polyproteins or as individual proteins. This occurs during the maturation process which transforms the inactive virion into a fully infective particle. During maturation, the viral RNA is aggregated and condensed by the different forms of the nucleocapsid which appear sequentially (39). A first cleavage occurs between the MA-CA-SP1 and the NC(p7)-SP2-p6 domains, thereby liberating transiently NC(p15) (*ie* NC(p7)-SP2-p6). The next cleavage releases the p6 domain from NC(p15) yielding the transient NC(p9) (*ie* NC(p7)+SP2). Finally, the last cleavage results in the liberation of the mature NC(p7)_1-55_ and SP2. On the contrary of mature NC(p7), NC(p15) and NC(p9) only appear transiently in the virion (70). Wang *et al.* (71) have compared the nucleic acids binding and chaperone efficiencies of these three forms *in vitro*. The authors deduced that the acidic residues in p6 modulate both the aggregation and annealing properties of NC(p15). Because the NA aggregation capabilities of NC(p15) are reduced compared to NC(p9) and NC(p7), and because the alanine substitution of acidic residues in p6 restores an efficiency comparable to NC(p9) and NC(p7), they suggested that p6 folds back on NC(p7) basic zinc fingers, establishing thus a competition with nucleic acids binding. Their assumption on an interaction between the NC(p7) and the p6 domains of NC(p15) is supported by the comparison of the [^1^H–^15^N] HSQC spectra of NC(p15) and NC(p7)_1-55_ recorded in water at 25mM NaCl, which evidences significant CSPs for residues F16, A25, K23 and K38, *ie* principally in the ZF1 region. This is consistent with our DSSP measurements in the presence of 10 eq. p6 and in DPC-*d_38_* (Fig. 5B), which demonstrate that the ZF1 region is preferentially targeted by p6 than ZF2. The originality of our results is that in our case p6 is not linked to NC(p7) and free to orient itself to interact.

The dissociation constant of the p6/NC(p7)_1-55_ complex we measured in DPC conditions, reflecting an interaction, falls within the hundred micromolar range. Such interaction may be stronger in the context of Gag, due to the increased likelihood of an intramolecular encounter between the two domains and to the interaction with the first zinc finger. It is therefore conceivable that a finely tuned regulatory mechanism exists among these players, with the p6 domain of Gag potentially engaging interactions in *cis* with NC(p7)_1-55_ or in *trans* with RNA, respectively influencing the assembly process, the dimerization of viral RNA and the multimerization of Gag. (Fig 9). Banerjee and *al*. (65) proposed that, once viral RNA is selected by the NC(p7) region, p6 regions from different Gag monomers could engage in stable inter-helical interactions through their respective C-terminal helix.

In conclusion, in the light of our results, HIV-1 Gag most probably exist as a mixture of open and closed forms of its C-terminal NC(p7)-SP2-p6 multidomain. The close form may bring protection to the poorly structured p6 towards the cellular proteases. The back-folded state of p6 could also regulate the binding of hosts factors like Tsg101 and ALIX or the recruitment of Vpr. Given the interaction observed between the free forms of NC(p7)_1-55_ and p6, we reasoned that these two cis-acting domains within Gag might function together to selectively recruit the viral gRNA. With its low affinity in aqueous buffer, but with enhanced susceptibility of interaction with NC(p7) in micellar conditions mimicking the vicinity of the plasmic membrane, p6 could be the conductor’s baton that orchestrate the selective recruitment of the gRNA. All these mechanisms need of course to be experimentally assayed.

## MATERIALS AND METHODS

### Protein expression and synthesis

^15^N labeled and unlabeled NC(p7)_1-55_ proteins were expressed and purified in *E. coli* as previously described (41, 72). ^15^N-NC(p7)_1-55_ NMR samples were prepared at a concentration around 0.4 mM, in a 25 mM sodium acetate-d4 buffer pH 6.5 containing 25 mM NaCl, 0.1 mM ZnCl_2_, 0.1 mM β mercaptoethanol and 10% ^2^H_2_O. Fully labeled DPC-*d_38_* was used for NMR experiment. Partially ^15^N-labeled p6^L^ and unlabeled p6 proteins were synthetised as previously described (27). Both for NMR and Fluorescence Anisotropy measurements, the DPC concentrations were set to a range of molar equivalents using the NC(p7) _1-55_ concentration as reference.

### NMR experiments

All NMR experiments were recorded at either at 10°C or at 30°C on a Bruker AVANCE III HD 600 MHz spectrometer equipped with a 5mm TCI cryoprobe with Z-gradients. Internal DSS standard was used for direct referencing of the ^1^H chemical shifts, and indirect referencing using gyromagnetic constants ratios for ^15^N and ^13^C chemical shifts (73). Chemical shifts of NC(p7)_1-55_ and p6 amino acids were measured on standard [^1^H-^15^N] 3D NOESY/TOCSY-HSQC (74, 75) and 2D NOESY. They have been deposited to BMRB under the accession number 35058 for NC(p7)_1-55_, 35060 for p6 in acetate buffer without DPC, and 35059 for p6 in 450eq. DPC-*d38*. Inter proton distances were computed from a 150 ms mixing time [^1^H-^15^N] 3D NOESY-HSQC. Titrations of the two proteins were followed by [^1^H-^15^N] SOFAST-HMQC (76). ^1^H-^15^N hetero-NOE experiments were measured with a recycling delay of 5s between two successive scans (77, 78).

Coupling constants ^3^J_HN-Hα_ and ^3^J_N-Hβ_ were derived respectively from 3D HNHA and HNHB experiments to yield ϕ (HN-N-Cα-Hα) and χ (N-Cα-Cβ-Hβ) angles using Karplus equations (79–81), which were introduced as constraints for structure calculations. The relaxation times experiments were recorded with a recycling delay of 5s. T1 data were collected on 14 experiments using values of 3, 20, 40, 60, 80, 100, 200, 300, 400, 500, 600, 700, 800, and 12000 ms as recovery delays. T2 experiments were collected on 12 experiments using CPMG lengths of 17, 34, 67, 84, 102, 135 (repeat), 168, 202, 204, 219, 238 ms with inter-pulse delays of 900μs between the ^15^N 180 degree pulses. All 2D and 3D experiments were processed with NMRpipe (82) and Sparky (83) softwares. Predictions of the backbone dihedral angles (Φ, Ψ) from the chemical shifts were carried out using TALOS software (84–86) and DANGLE in CcpNmr Analysis program (http://www.ccpn.ac.uk/software/analysis). The Secondary structure propensity (SSP) (87) scores were calculated using N, H^N^, Hα chemical shifts observed for free NC(p7)_1-55_ and with 10 eq. p6 in H_2_O in the absence and in the presence of 450 eq. DPC-*d_38_*. Chemical shift perturbations (CSPs) of the amide groups are computed according to Equation 1 (88) where Δδ^15^N and Δδ^1^HN are the chemical shift perturbations observed in the nitrogen and proton dimensions and α constant set to 0.2 for glycine and 0.14 for any other amino acid. We considered as strong the CSPs values that exceed 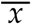 + *σ*, where 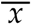 is the mean and σ the standard deviation. We excluded from these computations the CSPs of N55 measured in DPC in the presence of 0.5 and 10 eq. p6 (Fig. S6) which are greater than three times the standard deviation.

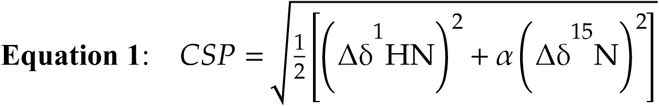

### NMR structures

Structure calculations were started from randomized structures of p6 or NC(p7)_1-55_ with two Zn atoms. A hexa-coordination angle of 109° and a partial charge delocalization on the 3 cysteines and the histidine were imposed on the two zinc fingers all along the simulated annealing (SA) and refinement procedures. SA calculations and refinement were carried with Xplor-NIH with the EEFx force field according to the protocols of *Tian et al*. (89, 90). NC(p7)_1-55_ structure calculation was restrained by 422 NOE-derived distances dispatched into strong [1.8Å-2.5Å], medium [2.5Å-3.5Å] and weak [4.5Å-6Å] distances, by 108 ^3^J_HN-Hα_ and 108 ^3^J_N-Hβ_ coupling constants and by the 98 ϕ and φ torsion angles predicted by TALOS from the chemical shifts. p6 structures were generated using 150 (H_2_O) or 307 (DPC) NOE distance restraints dispatched in the same strong, medium and weak distance intervals, and 80 or 98 restraints on ϕ and φ torsion angles obtained with TALOS software, respectively for 10°C in H_2_O and 30°C in DPC conditions.

NC(p15) starting 3D model was generated with AlfaFold3 (91). It was refined at 1500K against the same experimental restraints as for free p6 and NC(p7)1-55, using Xplor-NIH ans the EEFX force field. Procheck (92) and MolProbity (93) programs were used to validate the conformations of the models of structures. PDB accession numbers are as follows: 30ZZ for NC(p7)_1-55_, 31AD for p6 in water and 31AA for p6 in DPC. Structures were visualized using PyMOL 1.2 software.

### NMR titration of ^15^N-NC(p7)_1-55_ by p6

On [^1^H-^15^N] SOFAST-HMQC experiments recorded at 30°C, the variations of CSPs (Δ_obs_) with the concentration of p6 are followed. The reference experiment corresponds to the free NC(p7)_1-55_ protein. Δ_obs_variations were fitted to produce the apparent dissociation constant K_Δapp_ using the Equation 1 (94) representative of a 2-step model:

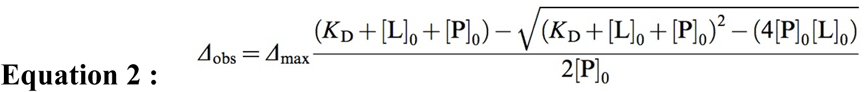

[P]_0_ is the reference concentration for free NC(p7) (100 μM). [L]_0_ is variable total concentration of p6 at each step of the titration. Δ_max_ is maximum CSP expected when all NC(p7)_1-55_ is complexed with p6.

### Fluorescence Anisotropy titration of NC(p7)_1-55_ by p6 with and without DPC

Fluorescence perpendicular and parallel polarized intensities were recorded at 23°C in a low-volume 384-well black flat bottom microplate on a Clariostar spectrometer (BMG Labtech) equipped with a dichroic filter at 325nm. Excitation and emission wavelengths were set respectively to 295 and 360nm with respective 10 and 20 nm bandwidths. Data were recorded with 200 flashes per point in orbital mode.

Two sets of samples were prepared with sample volumes of 7 and 10 μl. For each set, the NC(p7)_1-55_ concentration was kept constant to 10 and 1.3 μM respectively throughout the titration. Data points were recorded in duplicate after one-hour equilibration for increasing p6 concentrations corresponding to 0.5, 1, 5, 10, 50, 100, 500, 1000 eq. of NC(p7) _1-55_ concentration. The buffer for these assays was identical to the NMR buffer. Measurements were then carried out with increasing DPC concentrations (0, 5.5mM, 185mM, 548 mM, 886mM and 1.07M) using the two same sets of samples. Protein concentrations were corrected for the volume variations induced by DPC addition from a concentrated stock solution. Buffer contributions were removed by subtracting the intensities or the anisotropies of control buffer samples.

Fluorescence polarizations were computed from the perpendicular and parallel fluorescence intensities. Data were analyzed with Kaleidagraph software. Experimental polarizations F_obs_ were computed for each data point and compared to F_free_ of the free NC(p7)_1-55_. Titration data were fitted to produce the dissociation constant K_d_ using same Equation 1 as for NMR titrations, using a [P]_0_ concentration of either 1.3 or 10 μM depending on the assay. Δ_max_ corresponds to the fluorescence polarizations observed for full complex formation and is determined during the fitting procedure.

## ACKNOWLEDGMENTS

We thank Sarah Lagier Lopez, Juliette Lecomte and Xiaowei Chen for their help in protein NC(p7)_1-55_ expression. We wish to thank Serge Bouaziz and Rodrigue Marquant for providing the synthetic peptides p6 and p6^L^. We thank Pascale Coric for her help to access the NMR spectrometer. This work was supported by fundings of Université Paris Cité and CNRS. The authors declare to have no conflicting interests.

## Contributions

VL conceptualized and initiated the research. VL and SN-L designed all the experiments. VL ran all NMR experiments. SN-L ran all fluorescence experiments, produced, and purified a part of the recombinant NC(p7)_1-55_. VL and SN-L both ran molecular simulations. VL and SN-L both analyzed all the data, realized the figures and wrote the paper.

